# Method for Identifying Proteomic Biomarkers of Health

**DOI:** 10.1101/2023.12.06.570488

**Authors:** Ksenia Zlobina, Ashwin Gopinath, Anu Thubagere

## Abstract

Classification of proteomic samples from sick and non-sick individuals is important for developing high-quality diagnostics of diseases. Creating a shortlist of proteins useful for diagnostics is challenging. In this manuscript a simple algorithm of creating a multidimensional biomarker of health based on scanning proteomics data is provided. The algorithm is applied to several existing publicly available datasets and demonstrates a 9-protein indicator of atopic dermatitis, and 6-protein indicator of heart failure.

## INTRODUCTION

Disease prediction has always been an important problem in medicine. In recent decades, computer technologies came into the health sector, and big health-related datasets, such as medical history records, became available to researchers. This induced the development of algorithms for disease prediction based on big biological data [Yao 2013, McCormic 2011, Farran 2013, Ahmad 2013, Uddin 2019, Park 2021]. Proteomics (along with other -omics) provided additional health state measurements and has been actively explored in disease research [Zhong 2021, Gisby 2021, Brunner 2017, Chen 2020]. Proteomic measurements may help to find new indicators of diseases; however, it is not clear in advance how many of the provided measurements are informative.

Proteomics biomarkers of diseases are usually searched among highly differentially expressed values [Zhong 2021, Gisby 2021]. This usually leads to finding elevated or reduced levels of some proteins in sick people, but with highly overlapping protein values in sick and non-sick people, making high-quality diagnostics challenging.

The next step in searching for high-quality biomarkers is looking for multidimensional proteomic indicators. Researchers have used several methods to find an algorithm with a multidimensional proteomic vector as input and disease classification as output. These algorithms often involve neural networks or other complex algorithms [Gyllensten 2022].

A simplest multidimensional classification is dividing multidimensional space by a linear hyperplane. Linear classification of data must be applied in the space of several proteins, and the dimensionality (number of proteins defining the disease) of this space should not be too large. The dimensionality of the linear algorithm determines the number of fitting parameters. The number of data points that can be classified with a d-dimensional linear algorithm into any pair of classes is d+1 (in 2-dimensional space, any 3 points can be classified into 2 classes by a line).

This is explained for a wider range of classification algorithms by the concept of the Vapnik-Chervonenkis dimension [Vapnik 1971, Ischenko 2021]. Thus, the number of proteins defining linear classification must be smaller than the number of training data points, but for robust disease classification, it should be much smaller.

Selecting the shortlist of proteins suitable for high-quality linear classification of sick vs. non-sick sample data points is challenging [Gyllensten 2022].

In this manuscript, a method of finding a multidimensional biomarker of a health state is proposed. It implies scanning many possible subsets of proteins taken from a proteomics dataset and performing linear classification to find the best linear boundary between two classes of samples (from sick and non-sick patients). The method gives promising results on several publicly available proteomics datasets.

## THE ALGORITHM

Suppose that some proteomics dataset contains the values of N proteins in healthy and sick subjects.

### Step 1

> In the first step we test each pair of proteins: all healthy and sick subjects may be represented as points in the 2-dimensional space of two proteins. By applying linear discriminant analysis [Hastie, 2008], we can perform 2-class classification: find the best boundary between the regions where the majority of healthy subjects are, and the region with the majority of sick people. For each pair of proteins, there is some error of classification: number of wrongly classified subjects. At the end of the first step, the best pair of proteins is found: the pair that corresponds to minimal error. Selection of the first pair of proteins is the most time-consuming, as it requires application of the discrimination procedure (N*N/2)-N times.

### Step 2

> In the second step we apply the procedure of improvement of diagnostic proteins set by adding one more protein. This procedure is iterative, it may be applied as many times as needed.
>
> Suppose that the first k proteins have been selected and healthy/sick subjects are classified with minimal error. Consider (k+1)-dimensional set of proteins, consisting of k already selected and one new protein. Apply discrimination procedure of samples in (k+1)-dimensional space and estimate classification error. Find error for each new protein considered as a new dimension of (k+1) proteins.
>
> Select the protein with minimal error of (k+1)-dimensional classification. Add this protein to the selected set of proteins. The new set contains (k+1) protein, and the error is smaller or equal to the previous one.
>
> Redesignate (k+1) to k and repeat the second step. Repeat until the classification error becomes zero or sufficiently small.
>
> Each (k+1)-dimensional discrimination procedure is applied (N-k) times.

Thus, on each step, the algorithm increases the dimensionality diminishing the error of classification.

### Minimizing dimensionality

At any step of the algorithm, there may be a situation when more than one protein considered as (k+1)-th makes the error minimal. In this case, the best protein may be searched taking into account (k+2) and further proteins. This leads to the tree of selections.

Optimizing the tree of selection is out of the scope of this research. In this work, we consider (k+1)-th and then if needed, (k+2)-th protein; if the ambiguity still exists, the next protein is randomly selected from several proteins corresponding to the minimal error.

In the next section this algorithm is applied to the publicly available proteomic dataset from the blood serum of people with/without atopic dermatitis [Brunner 2017], and then to the heart failure dataset [Egerstedt, 2019].

## ALGORITHM APPLICATION

### Biomarkers of Atopic Dermatitis

Consider proteomic data presented in the supplementary material to the [Brunner 2017]. The authors compared proteomes of 61 atopic dermatitis patients, 19 psoriasis patients and 23 healthy people. Proteomics was measured in blood serum with the Olink test. Three Olink panels were used: Inflammatory, Cardiovascular II and Cardiovascular III. The authors provided supplementary with the tables of values of the most significant proteins in each sample: 75, 91 and 91 proteins from each panel correspondingly. In fact, this dataset is preliminary filtered by the authors by applying a specific procedure of finding the most relevant proteins.

We applied our algorithm to select proteins for diagnostics. The patients were divided into 2 classes: with atopic dermatitis (AD) and without atopic dermatitis (no AD), that means healthy and psoriasis patients are in the second class altogether. Selection of proteins was performed among 75 proteins of the Olink Inflammation Panel.

The first step of our algorithm results in a pair of proteins. Several pairs with equally minimal error were obtained. We selected one pair: Caspase 8 and SIRT2 proteins. Figure 1a shows how AD and no-AD samples are classified in 2-dimensional space (CASP8, SIRT2). Figure 1b shows the histograms of distribution of the samples by the values of the same proteins. It is seen that though SIRT2 is downregulated in AD compared to no-AD, there is an overlap between two classes that doesn’t allow to make good diagnostics. Representing the data in 2-d space (Fig 1a) makes diagnostics much better. Overall error, measured as the number of wrongly detected samples, in this case is 11, while the total number of samples is 103.

**Figure 1.**
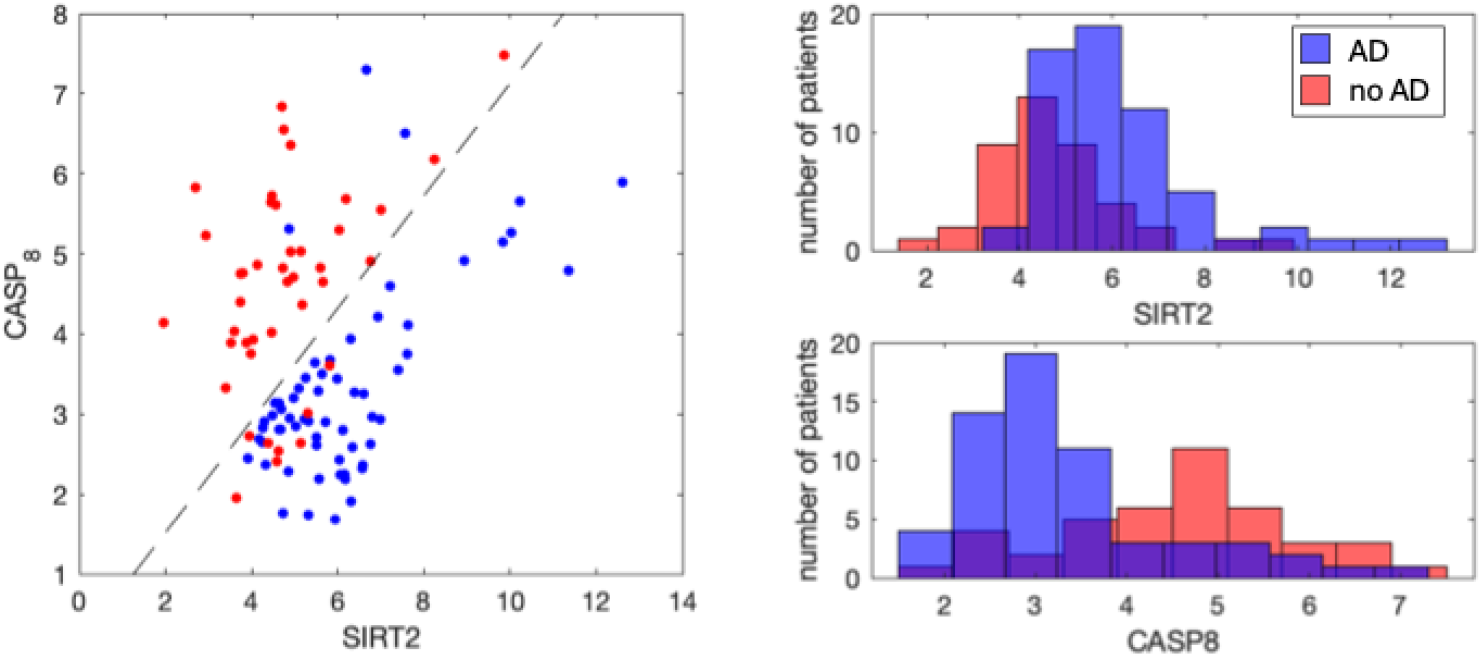
Representation of AD (blue) and no-AD (red) samples in 2-dimensional space of proteins (SIRT2, CASP8) - one of the pairs of proteins with minimal classification error (11 of 103 samples are wrongly detected). Histograms of sample distribution by each protein value are on the right.

Application of the second step of our algorithm results in consequent selection of proteins that increase dimensionality of the classification model and reduce the number of wrongly diagnosed samples. Figure 2 shows a number of wrong predictions for each number of proteins in the model.

**Figure 2.**
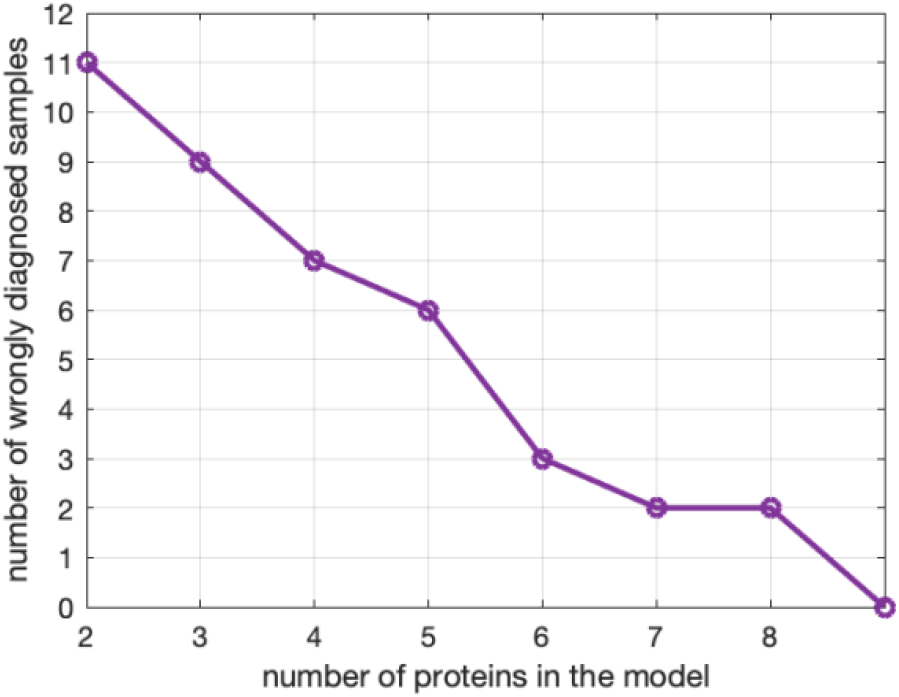
Number of wrongly diagnosed samples (error) after each step of the algorithm. Beginning from (SIRT2, CASP8) pair of proteins with error 11 of 103, the algorithm selects one more protein and calculates error. (See list of the proteins in table 1).

One can see that for the given run of the algorithm, 9 proteins are enough to clearly separate all data samples in 9-dimensional protein space. 9 proteins selected for diagnostics are presented in table 1.

**Table 1.** Linear coefficients for the detection of AD-index.

| Protein | bi | Protein | bi | $a$ |
| --- | --- | --- | --- | --- |
| SIRT2 | 1.03557989 | STAMBP | -3.657224 | -13.050641 |
| CASP_8 | 2.92219465 | TGF_alpha | -0.7417068 |  |
| EN_RAGE | -2.7527468 | CCL28 | -0.8827546 |  |
| IL_20RA | 1.55613736 | TWEAK | 2.11921052 |  |
| CD5 | 2.35761831 |  |  |  |

Figure 3 shows histograms of data samples distribution by protein value for each of these 9 proteins. It is seen that being taken separately no one protein can be a biomarker of atopic dermatitis, as there is huge overlap of AD and non-AD samples in each histogram. However, taken together as a multidimensional biomarker, these proteins can help to separate AD from non-AD using linear combination.

**Figure 3.**
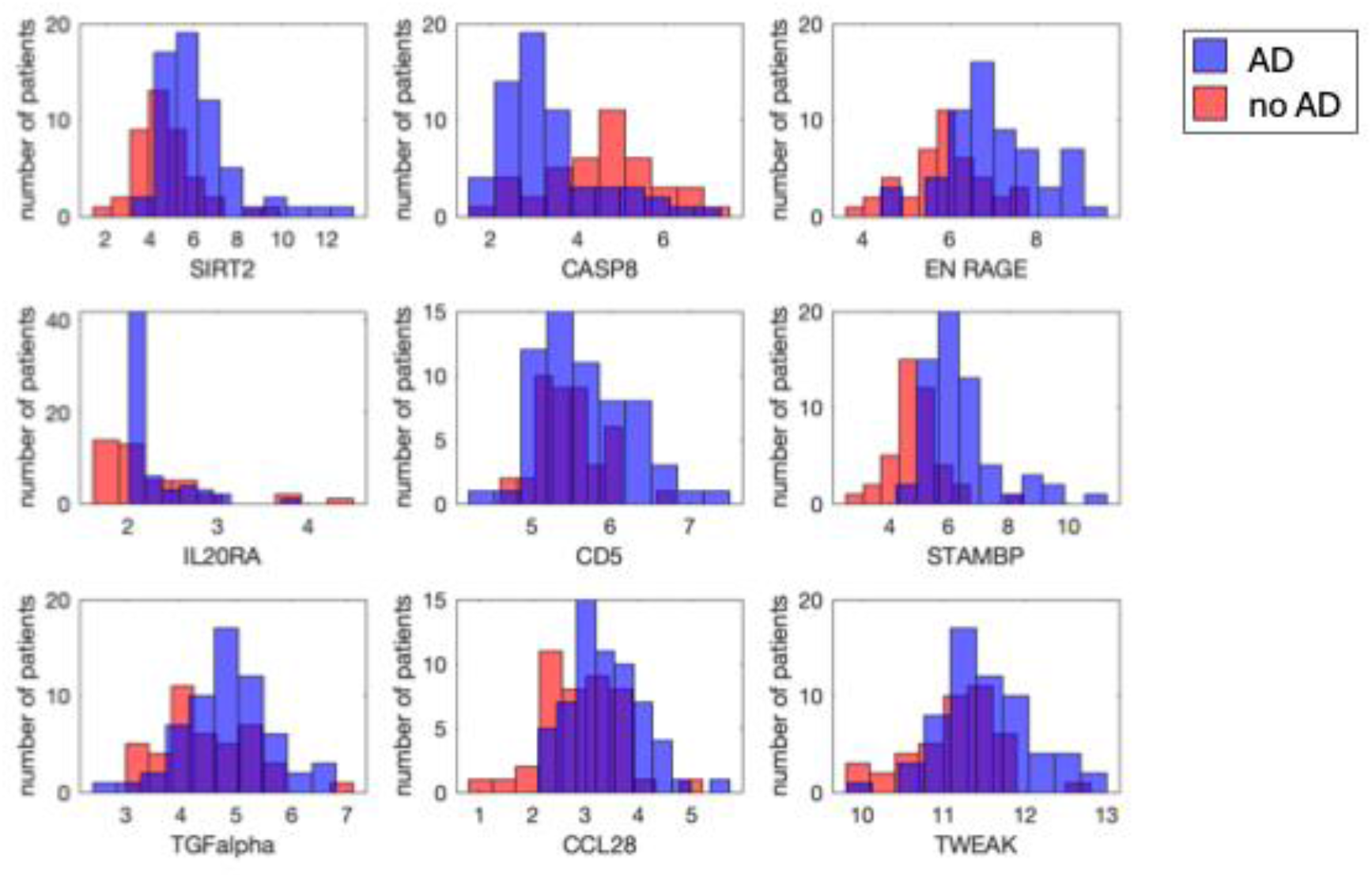
Histograms of sample distribution by the values of the protein from the set selected by algorithm for multidimensional biomarker. Blue: AD, red: no-AD patients.

**Figure 4.**
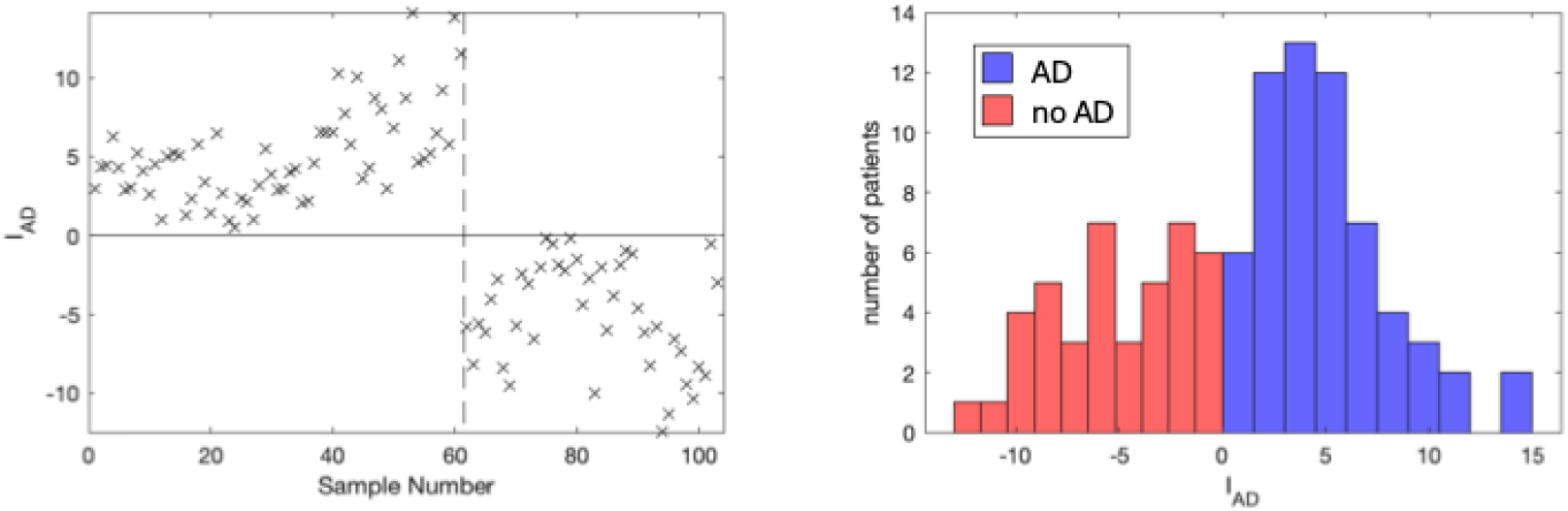
Left: Atopic dermatitis index based on 9 proteins for each patient. Samples 1-61 are from atopic dermatitis patients, samples 62-103 are from healthy or non-AD sick people. Right: histogram of distribution of patients by the value of index I_AD_.

To classify the samples one can use the index of atopic dermatitis *I*_*AD*_: a linear combination of 9 proteins:

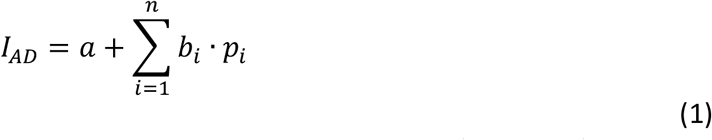

with n=9, where *p*_*i*_ is the proteomics value corresponding to protein *i*(*i* = 1 …*n*) and *b*_*i*_ is corresponding coefficient represented in table 1 for each protein. Coefficient *a* makes *I*_*AD*_ positive for AD samples and negative for those taken from no-AD subjects. The values of *I*_*AD*_ for all samples in the dataset are shown in Fig. 5.

**Figure 5.**
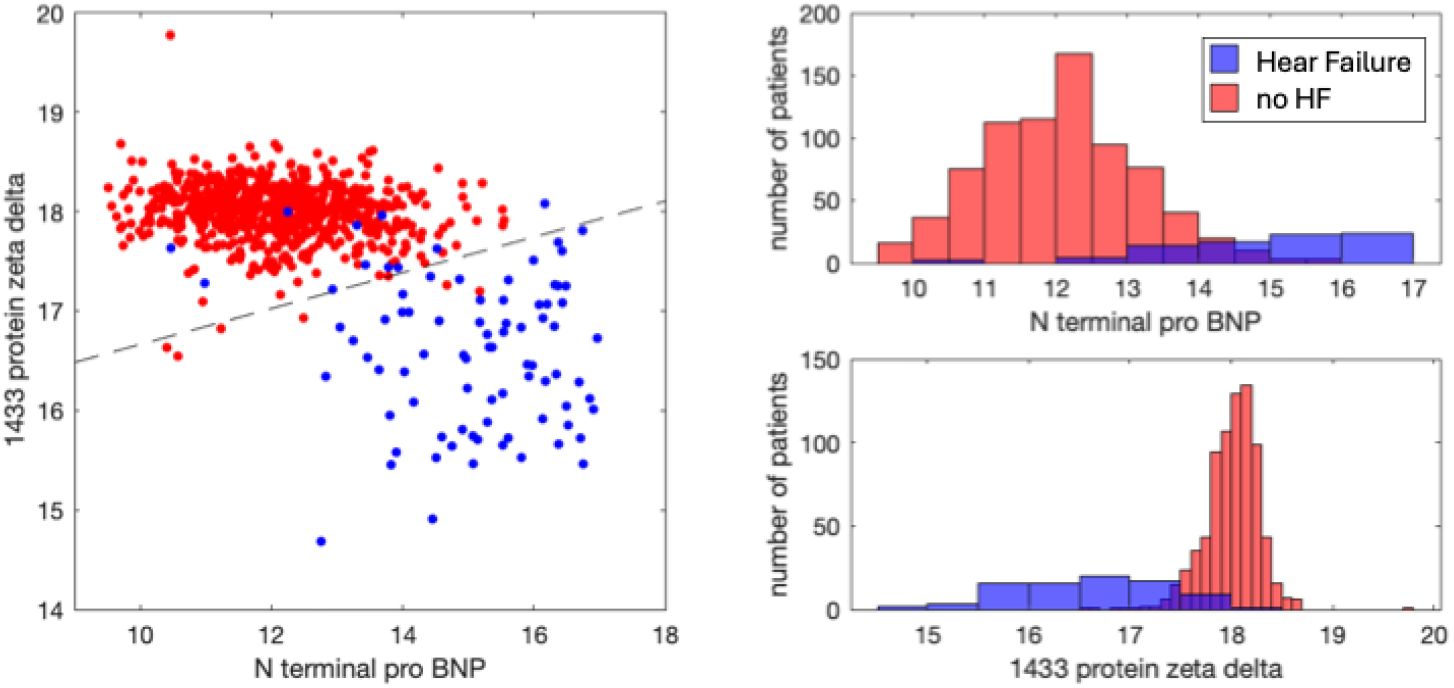
Representation of samples from patients with manifest heart failure (blue) and no heart failure (red) in 2-dimensional space of proteins (N terminal pro BNP, 1433 protein zeta delta) – a pair of proteins with one of the smallest classification errors (17 of total 852 samples are wrongly detected). Histograms of sample distribution by each protein value are on the right.

We argue that at least several sets of proteins and corresponding linear combinations can be selected for high-quality classification and diagnostics. Additional set of proteins for AD diagnostics is presented in Supplementary material S1.

### Biomarkers of heart failure

We test our algorithm for finding proteins related to heart failure using data provided in supplementary material in [Egerstedt, 2019]. This dataset has not been preliminary filtered by the authors, it contains information about 1305 proteins detected by SomaScan assay. There are several cohorts of patients of which we select two: 768 patients without heart failure and 84 patients with manifest heart failure. We use log2 of the proteomics values represented in the initial dataset. (The protein PBEF is a very good candidate to biomarker by itself. To make our mathematical problem more challenging, PBEF was removed from the dataset before application of the algorithm).

Several pairs of proteins may be selected after the application of the first step of our algorithm. We select the pair (N terminal pro BNP, 1433 protein zeta delta) as the first pair of proteins. The distribution of data samples by these two proteins is shown in Figure 5.

The second step of the algorithm quickly converges to error 0: 6-proteins are enough to classify healthy vs heart failure manifest patient samples (Figure 6).

**Figure 6.**
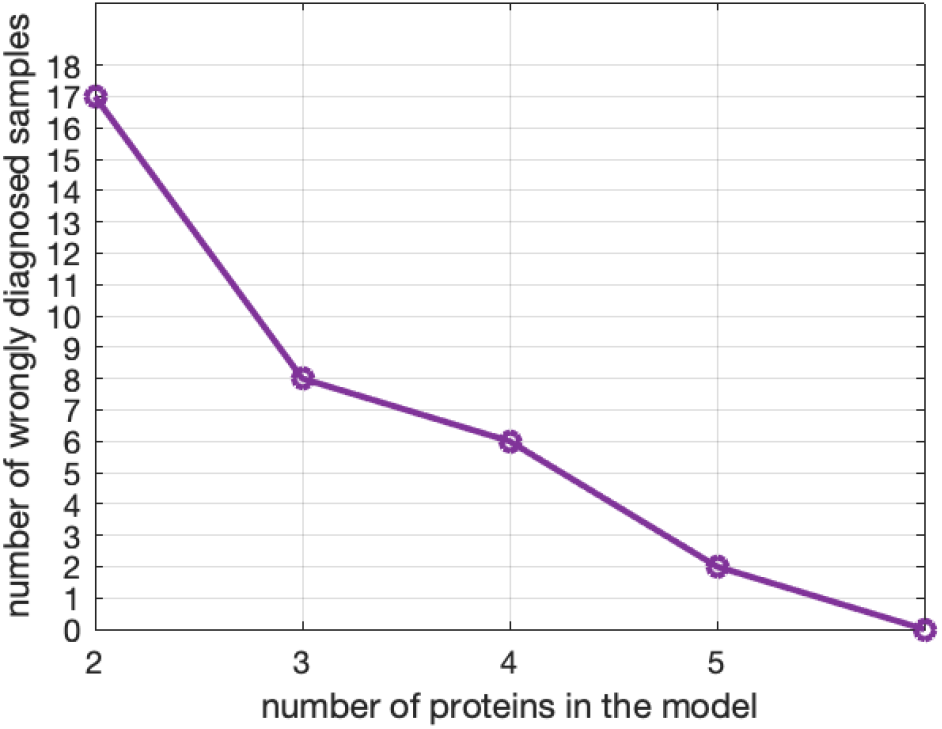
Number of wrongly diagnosed samples (error) after each step of the algorithm. Beginning from (N terminal pro BNP, 1433 protein zeta delta) pair of proteins with error 17 of 852, the algorithm selects one more protein at each step and calculates error.

Using a 6-protein model we created a proteomics index of heart failure I_HF_ (see Figure 7). For calculation, proteins represented in table 2 with corresponding coefficients must be substituted to formula (1).

**Figure 7.**
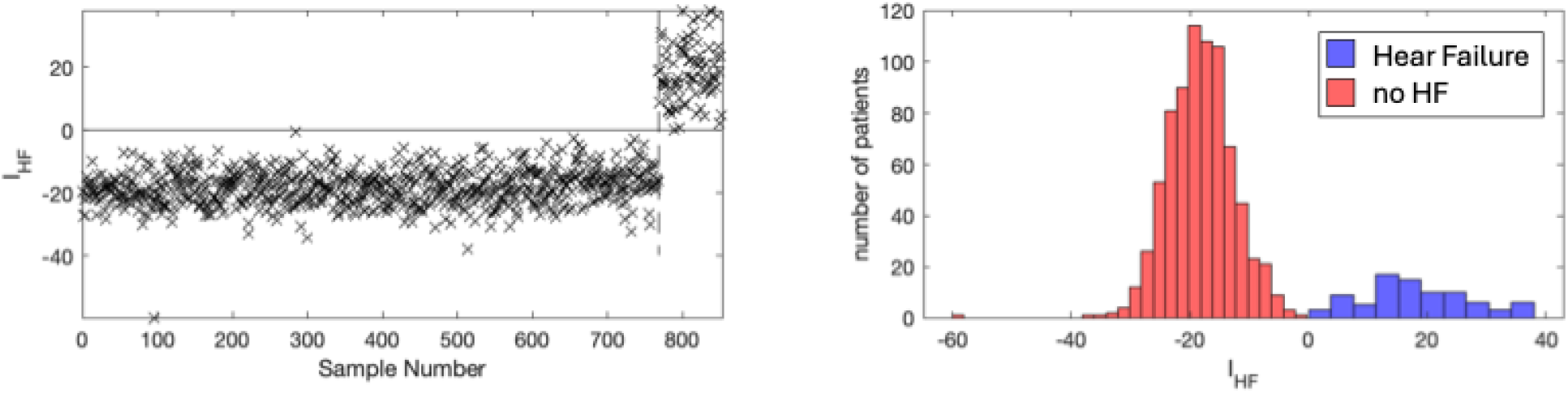
Heart failure index based on 6 proteins for each patient. (A) Samples 1-768 are from non-heart failure patients, samples 769-852 are from people with manifest heart failure. (B) histograms of distribution of HF-manifest samples (blue) and non-HF samples (red) by index of heart failure.

**Table 2.** Linear coefficients for the detection of heart failure index.

| Protein | bi | Protein | bi | a |
| --- | --- | --- | --- | --- |
| NterminalproBNP | -2.4335558 | H31 | 5.19069013 | -96.389203 |
| A_1433proteinzetadelta | 12.8852434 | IGFBP1 | 1.82859041 |  |
| SHC1 | -5.8183962 | DSC2 | -8.1055311 |  |

Figure 8 shows the distribution of samples by the values of each protein making contribution to the 6-protein classification model derived by algorithm. It is seen that no one protein can separate HF from non-HF samples by itself. But together they give rise to a good linear classification (Fig.7).

**Figure 8.**
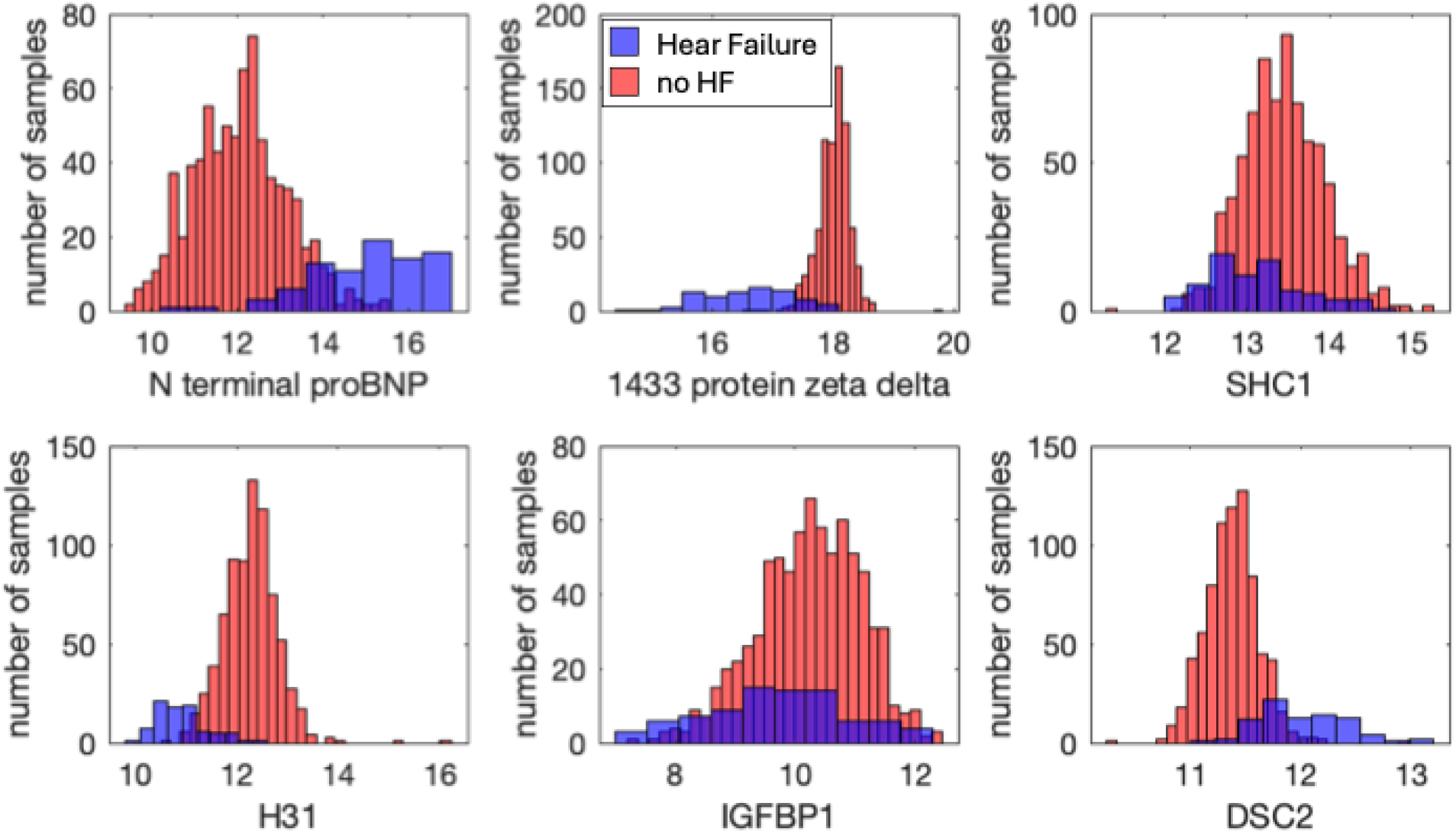
Histograms of sample distribution by the values of the protein from the set selected by algorithm for multidimensional biomarker. Blue: samples from heart failure, red: from non-HF.

## DISCUSSION

Proteomic analysis of a bio-sample reflects the levels of many proteins participating in many processes. A general goal of any proteomic research is to find information significant in the considered disease in this big data. Many proteomic measurements should be omitted as irrelevant to the considered biological process.

Diseases are dynamic biological processes, where at least some variables change in time. Proteomic variables collected from disease samples can be divided into three categories: (i) dynamic variables, (ii) constant values that determine the background of the ongoing process, and (iii) irrelevant variables - time-dependent or constant - but poorly related to the considered disease.

The <u>dynamic variables</u> of the disease are of greatest interest for understanding the nature of the disease. However, suitable temporal resolution of proteomic sample collection is required. Accurate mapping of multiomics sample collection time to the stage of the considered disease may be challenging as well [Filbin 2021].

Some proteins stay <u>constant</u> at all stages of the disease and provide the background environment for the disease. These proteins should be reliably up- or downregulated in all sick versus non-sick samples. For example, it was found that IL-13, CCL17, eotaxin-1/CCL11 are increased in atopic dermatitis [Brunner 2017], Nt-proBNP, CRP, ST2, troponin T, and TNFα were found to be higher in subjects with incident of heart failure than without it [Egerstedt 2019].

Even before the era of big data, biological systems were investigated by mathematical modeling [Murray, 1989]. One can imagine that proteomic data is measurements from some mathematical model simulations. Equations of a mathematical model consist of <u>dynamical variables</u> and <u>constant parameters</u>; both define transition of the biological system from health to disease. (Irrelevant variables are usually not considered in modeling but appear in proteomic data).

Dynamical variables should be harder to detect from proteomic data as they vary at different stages of the disease. Constant parameters are defined by levels of some proteins that stay constant during the biological process. Levels of these proteins can switch all the system from disease to health. They should be easier to detect in proteomic data analysis.

Parametric diagrams showing how constant parameters switch the biological system from one type of behavior to another have been found by mathematical modelling for many diseases [Domínguez-Hüttinger 2016, Arias 2015, Kartukha 2006, Zlobina 2016]. Analysis of proteomics is another way of finding parametric diagrams, without modeling, directly from data. Maybe in the future we will be able to map findings from mathematical models to data-derived results.

In recent years bioinformatic analyzing proteomic data is aimed at finding hidden relationships between proteins-participants of unknown biological networks [Chen 2020, Brunner 2017, Filbin 2021, Gisby 2021]. This is a very difficult task: in fact, the biological system is a black box, and the researchers try to learn it by analyzing the output signal only.

The method presented in this manuscript allows creating a multidimensional biomarker of a disease based on few proteins. Optimization procedure is needed to find the shortest list of proteins. Our tests with the model show that the list of proteins that can provide high-quality classification is not unique. For example, one more set of 8 proteins suitable for atopic dermatitis diagnostics is presented in the Supplementary material S1.

Similar observation was obtained in [Gyllensten 2022], where the authors used another approach and found that several different sets of proteins may classify ovarian cancer vs healthy patients’ samples. This may be because in a biological process some group of proteins may have similar dynamics, so in the model for diagnostics one protein can replace another (see example in Supplementary S2).

For better predictive ability of the model, non-correlated biomarkers are more useful, because two correlated biomarkers reflect the same information about the system, and one biomarker doesn’t add new information to another [Wang 2011]. Traditionally, new biomarkers of the diseases are invented based on known mechanisms of the disease and often correlate with existing biomarkers. High-throughput proteomic research may help to find non-obvious biomarkers, representing the unknown biological pathways [Wang 2011]. On the other hand, proteomics allows to find clusters of correlated proteins, so cluster protein levels could be used instead of single proteins.

Gyllensten and co-authors [Gyllensten 2022] note that many of the proteins included in their multivariate ovarian cancer models do not reach univariate statistical significance and they have not been previously detected in literature as being connected to ovarian cancer in the existing literature. Similar situation appears in our analysis: a single protein from the classification model cannot be used for classification (see histograms on the right in Figures 1 and 5). But being taken together with other proteins they contribute to better classification. Many proteins selected by our algorithm for diagnostics of atopic dermatitis or heart failure are not known to be related to corresponding health states.

Several modifications of this algorithm might improve it. For example, instead of a linear boundary between sick/non-sick samples, nonlinear discriminant analysis may be applied; or, instead of discriminant analysis, the boundary may be found based on the support vector machine (SVM) method. Whereas discriminant analysis looks at all points in each class and assumes a Gaussian distribution for each class, the support vector machine considers the edge points only [Hastie 2008, McLachlan 2004]. The number of these points depends on the settings, that may be varied to find better classification.

The algorithm developed in this work is easy to apply for diagnostics: one can take a table of coefficients for each protein, patient proteomics data and substitute them into formula (1). The model gives an answer to the “yes-no” question, i.e. the algorithm is suitable for bi-class classification. Future question is how to include it into multi-class models? What if overlapping lists of proteins detect two different diseases, how to define if the patient has disease A rather than disease B?

The experimental way how the proteomics analysis is taken is important. The resulting formulas with specific coefficient revealed here may be applied to another proteomic analysis only if it has been done with the same experimental platform (in our case: Olink for AD-classification and SomaLogic for HF-classification), with the same biomaterial (serum, blood, or plasma). Transition from one experimental platform to another or from one tissue to another must be done with care, additional experiments and special calculations are needed [Haslam 2022, Kim 2018].

The presented algorithm is directed to select a set of proteins for linear multidimensional bi-class classification of health state. Optimization is needed as well as building hierarchical structure for multi-class classification. If we will resolve all these problems, probably, we will become a bit closer to the diagnostics of health by one droplet of blood.

## SUPPLEMENTARY MATERIAL

### S1. Another set of proteins for atopic dermatitis prediction

There are several sets of proteins suitable for AD/no-AD diagnostics, and here we show one of them. If after the first step of the algorithm we select the pair (STAMBP, CASP8), our algorithm comes to 8 proteins for classification without errors on this dataset (Figures S1 and S2).

Corresponding index of atopic dermatitis IAD can be calculated using linear formula (1) with coefficients represented in Table S1. Figure S3 shows the values of this index for all samples of the dataset.

**Figure S1.**
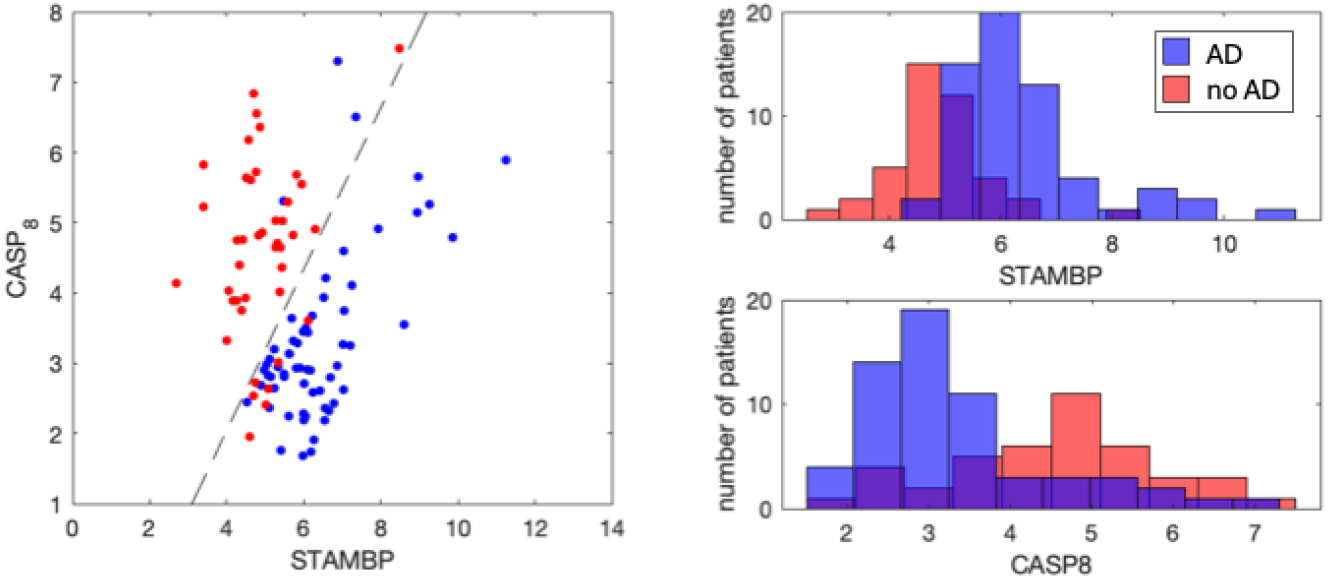
Representation of AD (blue) and no-AD (red) samples in 2-dimensional space of proteins (STAMBP, CASP8) - one more pair of proteins with minimal classification error (11 of 103 samples are wrongly detected). Histograms of samples distribution by each protein value are on the right

**Figure S2.**
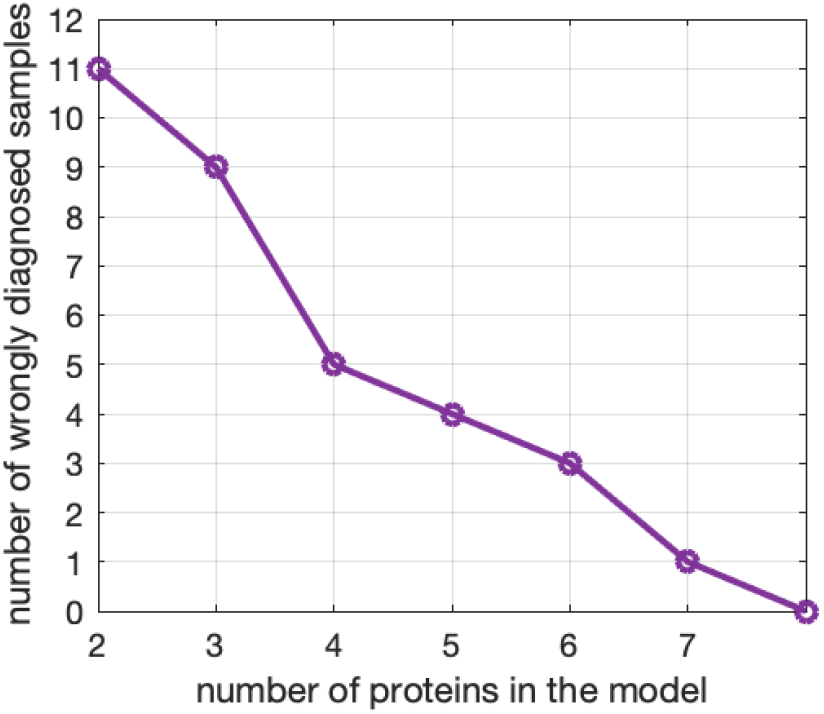
Number of wrongly diagnosed samples (error) after each step of the algorithm. Beginning from (STAMBP, CASP8) pair of proteins with error 11 of 103, the algorithm selects one more protein and calculates error. (See list of the proteins in table S1).

**Figure S3.**
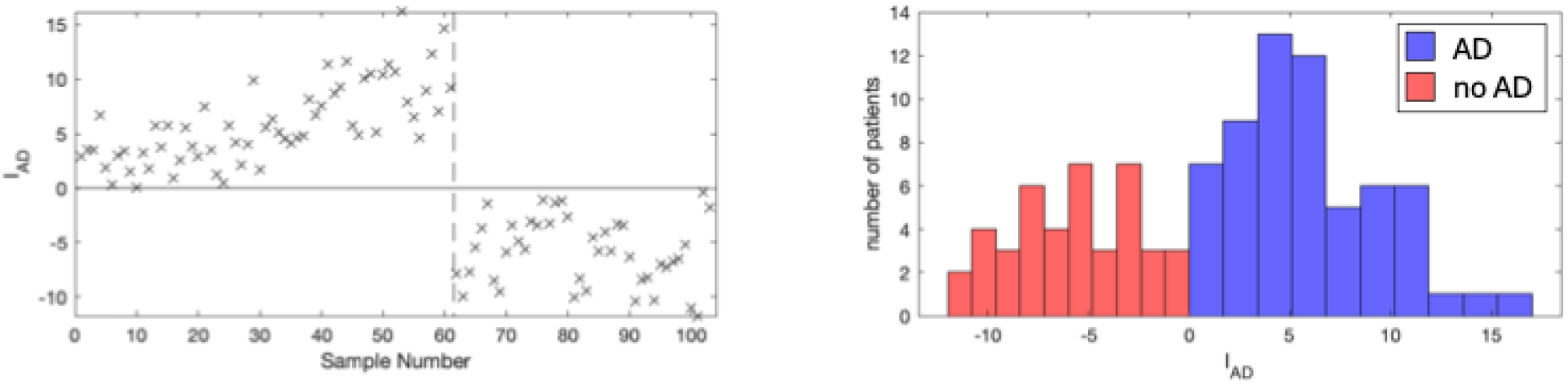
Left: Atopic dermatitis index based on 8 proteins. Samples 1-61 are from atopic dermatitis patients, samples 62-103 are from healthy or non-AD sick people. Right: histogram of distribution of patients by the value of index I_AD_.

**Table S1.**
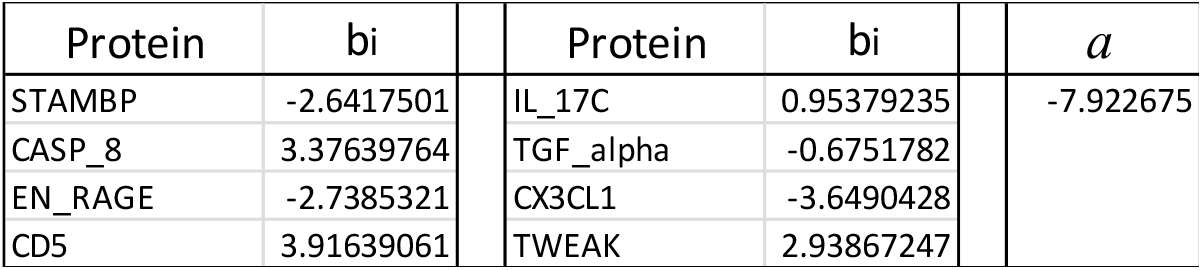
Linear coefficients for the detection of AD-index.

| Protein | bi | Protein | bi | $a$ |
| --- | --- | --- | --- | --- |
| STAMBP | -2.6417501 | IL_17C | 0.95379235 | -7.922675 |
| CASP_8 | 3.37639764 | TGF_alpha | -0.6751782 |  |
| EN_RAGE | -2.7385321 | CX3CL1 | -3.6490428 |  |
| CD5 | 3.91639061 | TWEAK | 2.93867247 |  |

**Figure S4.**
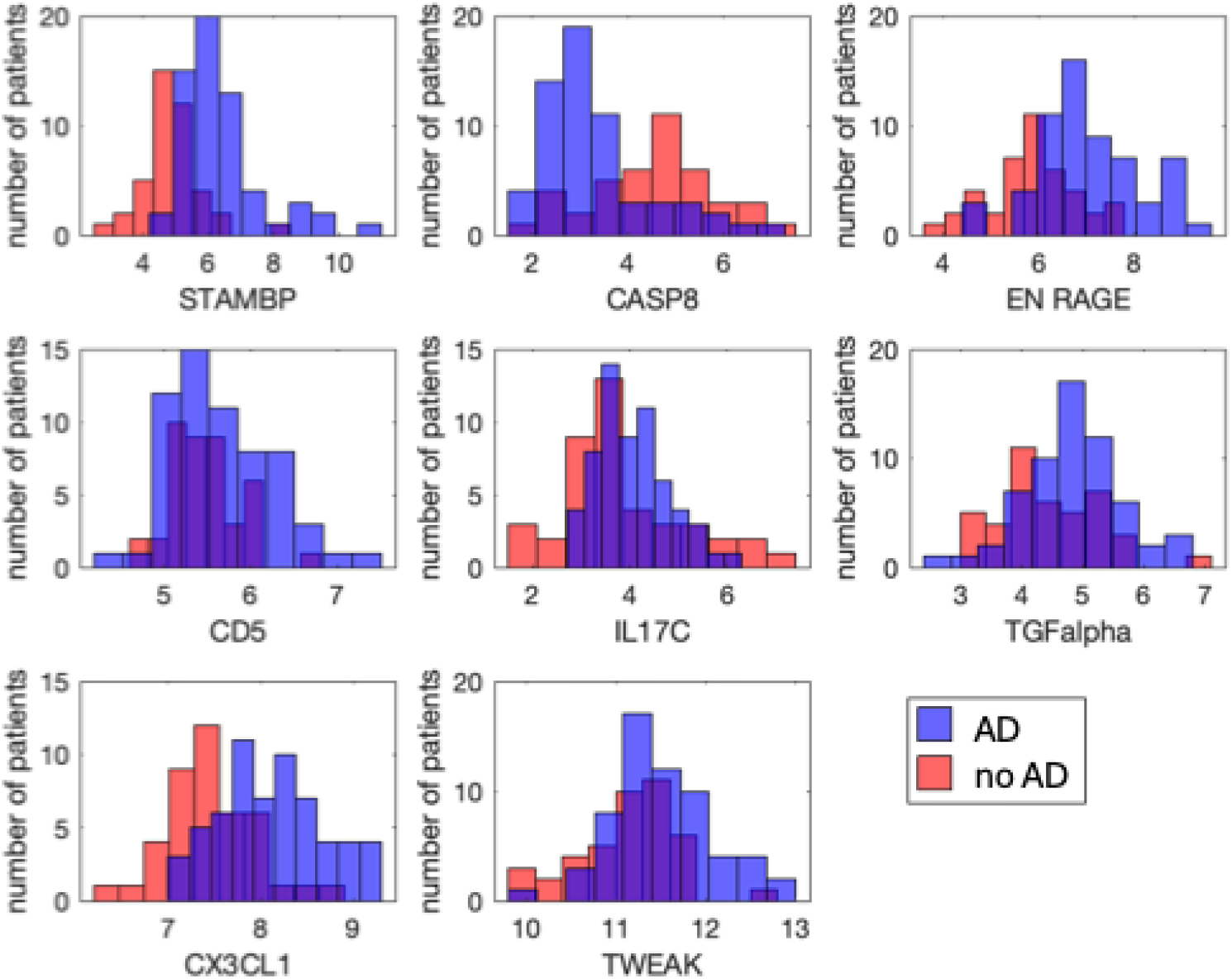
Histograms of samples distribution by the values of the protein from the set selected by algorithm for multidimensional biomarker. Blue: AD, red: no-AD patients.

Figure S4 demonstrates the histograms of distribution AD and non-AD samples by level of each protein. No one of the proteins can be a good biomarker but taken together as one multidimensional biomarker these proteins can be used for diagnostics.

Thus, the combination of proteins suitable for classification of sick vs non-sick samples is not unique.

### S2. Replacing protein in classification model by another, correlated protein

Some proteins in the dataset may be highly correlated with others. This means, that the relation between the values of two proteins in the same sample may be approximated by linear formula:

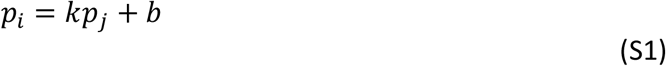

where *p*_*i*_ and *p*_*j*_ are the values of two proteins in the same sample; *k* and *b* are coefficients. This means that in the model of linear classification the protein *p*_*i*_ may be replaced by linear combination given by formula (S1).

Consider the classification model shown is supplementary S1. We first search proteins highly correlated with the proteins of the model. For proteins STAMBP, CD5 and CX3CL it is possible to find other proteins, with correlation coefficients above 0.8 (Table S2).

An example of plot of the proteins from the classification model (STAMBP) versus another protein (SIRT2) with correlation coefficient 0.93 is shown in figure S5. Linear formula for approximation for this data is STAMBP=0.72*SIRT2+1.79.

Coming back to calculation of AD-index (table S1 and formula (1)), we replace STAMBP values by 0.72*SIRT2+1.79. The obtained index of atopic dermatitis I_AD_ can classify AD and non-AD samples very well. The error of classification is 2 wrongly detected samples of 103.

The results of replacing one protein by linear approximation by another protein in the classification model are represented in Table S2.

**Table S2.** Replacing protein by another highly correlated protein in the classification model. The first protein in each pair was selected from the set derived in S1 supplementary. The protein in the classification model may be replaced by linear approximation using another protein. Corresponding error of classification (number of wrongly classified samples) is shown in the last column.

| Pair of proteins |  | Correlation coefficient | Linear approximation | Classification error |
| --- | --- | --- | --- | --- |
| STAMBP | SIRT2 | 0.933547 | $STAMBP = 0.71610552 * SIRT2 + 1.78646247$ | 2 |
| STAMBP | ADA | 0.827738 | $STAMBP = 0.99782888 * ADA - 1.0199372$ | 8 |
| STAMBP | x4E_BP1 | 0.848634 | $STAMBP = 0.53201975 * x4E\_BP1 + 2.38653357$ | 3 |
| CD5 | TNFRSF9 | 0.878702 | $CD5 = 0.69146217 * TNFRSF9 + 0.16129386$ | 3 |
| CX3CL1 | CSF_1 | 0.806 | $CX3CL1 = 1.24480965 * CSF\_1 - 4.194792$ | 3 |

**Figure S5.**
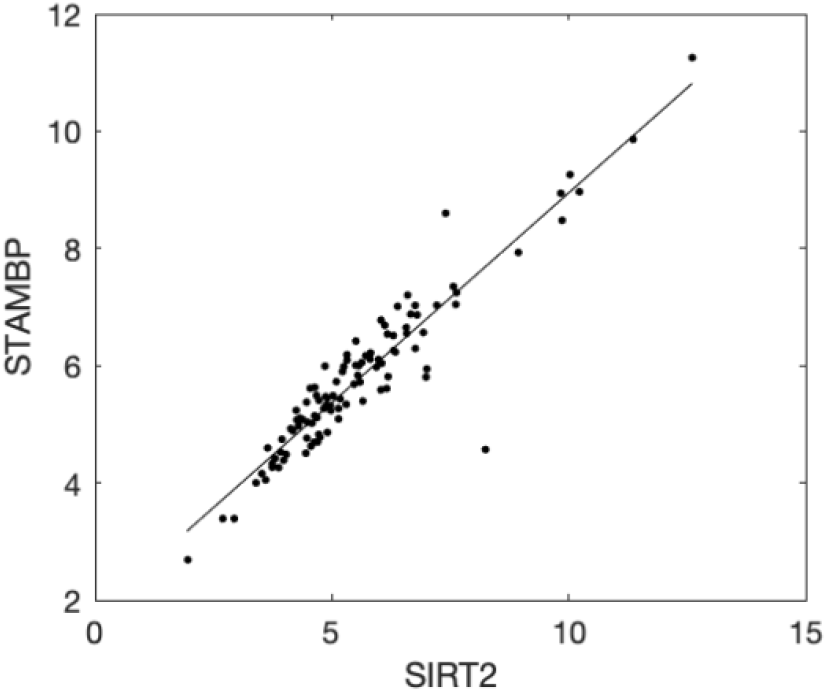
STAMBP and SIRT2 proteins levels in the samples from the atopic dermatitis dataset. One point represents one sample. Because of high correlation between two proteins, one protein can replace another in the health classification models. STAMBP protein values may be approximated by linear formula STAMBP=0.72*SIRT2+1.79.

